# Phospho-sRNA-seq reveals extracellular mRNA/lncRNA fragments as potential biomarkers in human plasma

**DOI:** 10.1101/553438

**Authors:** Maria D. Giraldez, Ryan M. Spengler, Alton Etheridge, Annika Jane Goicochea, Missy Tuck, Sung Won Choi, David J. Galas, Muneesh Tewari

## Abstract

Extracellular RNAs (exRNAs) in biofluids have attracted great interest as potential biomarkers. Whereas extracellular microRNAs (miRNAs) in blood plasma are extensively characterized, extracellular messenger RNAs (mRNA) and long noncoding RNAs (lncRNA) are less well-studied. We report that plasma contains fragmented mRNAs and lncRNAs that are largely missed by standard small RNA-seq protocols due to lack of 5’ phosphate or presence of 3’ phosphate. These fragments were revealed using a modified protocol (“phospho-sRNA-seq”) incorporating RNA treatment with T4-polynucleotide kinase, which we compared with standard small RNA-seq for sequencing synthetic RNAs with varied 5’ and 3’ ends, as well as human plasma exRNA. Analyzing phospho-sRNA-seq data using a custom, high-stringency bioinformatic pipeline, we identified mRNA/lncRNA transcriptome fingerprints in plasma, including tissue-specific gene sets. In a longitudinal study of bone marrow transplant patients, bone marrow-and liver-enriched exRNA genes tracked with bone marrow recovery and liver injury, respectively, providing proof-of-concept validation as a biomarker approach. By enabling access to an unexplored realm of mRNA and lncRNA fragments, phospho-sRNA-seq opens up new possibilities for plasma transcriptomic biomarker development.

## Introduction

In recent years, the discovery of a variety of extracellular RNA (exRNA) molecules present in the human bloodstream and other biofluids has been of great interest given their potential value as minimally-invasive biomarkers for a wide range of diseases(Freedman *et al*, 2016; Max *et al*, 2018; Godoy *et al*, 2018; Yuan *et al*, 2016). To date, characterization of exRNAs in blood has mostly focused on miRNAs, which have been shown to be exceptionally stable in plasma (i.e., the acellular portion of blood) by virtue of being protected in complexes with Argonaute proteins and extracellular vesicles(Arroyo *et al*, 2011; Hunter *et al*, 2008). However, miRNAs represent a small fraction of the human transcriptome and only a small minority of miRNAs show exquisite tissue- or disease-specificity(Ludwig *et al*, 2016). The degree to which the more predominant components of the transcriptome, notably mRNAs and lncRNAs, are similarly represented in blood as exRNA is less well established. Yet mRNAs and lncRNAs are highly appealing from the standpoint of biomarkers for monitoring health and disease due to their multiple established tissue- and disease-specific gene expression signatures(Perou *et al*, 2000; Potti *et al*, 2006; Chen *et al*, 2007; Ben-Porath *et al*, 2008; Iyer *et al*, 2015; Liu *et al*, 2008a).

RNA-seq has transformed transcriptome characterization in a wide range of biological contexts(Mortazavi *et al*, 2008; Wang *et al*, 2009) including its application to analyze exRNA in body fluids(Adiconis *et al*, 2013; Giraldez *et al*, 2018). These efforts have begun to elucidate the complex composition of exRNA in blood(Freedman *et al*, 2016; Max *et al*, 2018; Yeri *et al*, 2017; Godoy *et al*, 2018). There have been indications of extracellular mRNA and lncRNA in some studies of plasma, but results have been inconsistent, with some profiling studies reporting a variable percentage of them and others not even reporting their presence(Freedman *et al*, 2016; Max *et al*, 2018; Godoy *et al*, 2018; Danielson *et al*, 2017; Koh *et al*, 2014; Yuan *et al*, 2016; Yeri *et al*, 2017; Huang *et al*, 2013). Moreover, these profiling studies have used a variety of methods to evaluate exRNA expression (e.g. microarrays and different methodologies for RNA-seq) which, not surprisingly, contributes to the variation in findings across the studies.

We hypothesized that given the high concentration of RNases in the human bloodstream(Kamm & Smith, 1972), mRNAs and lncRNAs, if truly stable in blood plasma at all, may not exist in full-length form, but rather as small fragments. Furthermore, we hypothesized that standard ligation-based small RNA-seq methods might not detect such fragments because they are designed to capture miRNAs(Hafner *et al*, 2008), which by virtue of being products of RNase III class enzymes (e.g., Dicer) consistently present 5′-monophosphate and 3′-hydroxyl ends(Lee *et al*, 2003). In contrast, the 5′ and 3′ ends of RNA cleavage products generated by other ribonucleases vary substantially, which might prevent efficient adapter ligation with typical small RNA-seq methods. For example, abundant RNases in human blood circulation, such as those belonging to the ribonuclease A superfamily(Lu *et al*, 2018) degrade RNA dinucleotide bonds, leaving a 5′ hydroxyl and 3′ phosphorylated product (Cuchillo *et al*, 2011). Therefore, we reasoned that in order to comprehensively sequence a broader space of exRNAs beyond miRNAs, it would be essential to develop modifications to small RNA-seq protocols that can enable capture of RNA fragments that may have these alternate 5’ and 3’ phosphorylation states.

Here, we modified the standard small RNA-seq approach by incorporating both an upfront 5’ RNA phosphorylation / 3’ dephosphorylation step using T4 polynucleotide kinase (referred to here subsequently as “PNK”) and a custom, high-stringency bioinformatic data analysis pipeline to analyze non-miRNA small RNA fragments. This approach, which we refer to as “phospho-sRNA-seq”, revealed a large, untapped space of mRNAs and lncRNA fragments present in plasma. These fragments comprised tissue-specific signatures that were able to reflect biological processes of bone marrow reconstitution and acute liver injury in bone marrow transplant patients. We propose that this approach opens up new opportunities for disease biomarker discovery through transcriptomic analysis of exRNA fragments in the circulation.

## Results

### Synthetic RNA-based technical validation of a phospho-sRNA-seq protocol for recovering short mRNA and lncRNA fragments with ends lacking a 5’-phosphate and/or possessing a 3’-phosphate

To evaluate the performance of both standard and phospho-sRNA-seq methods for recovering short oligonucleotides with varying end-modifications likely to be found in human biofluids, we designed a synthetic reference pool comprising 476 ribonucleotides of different length (from 15 nt to 90 nt) and sequence (***Appendix Table S1***). More specifically, our pool includes 286 human miRNAs, 8 plant miRNAs, 164 fragments of mRNA and lncRNAs ranging from 15 nt to 90 nt and including different end modifications (i.e. 5’ phosphorylation, 3’ phosphorylation and none modifications) and 18 artificial miRNA sequences as control. As depicted in ***Figure 1A***, we prepared small-RNA libraries using this pool as input and following two different strategies: (i) standard ligation-based methodology (i.e. TruSeq small RNA protocol) and (ii) our modified approach of phospho-sRNA-seq (i.e. RNA pretreatment with PNK which phosphorylates 5’ hydroxyl groups and removes 3’ phosphoryl groups from oligonucleotides, followed by standard small RNA library preparation methodology). Libraries were multiplexed, sequenced on a NextSeq platform and analyzed as described in Methods.

**Figure 1.**
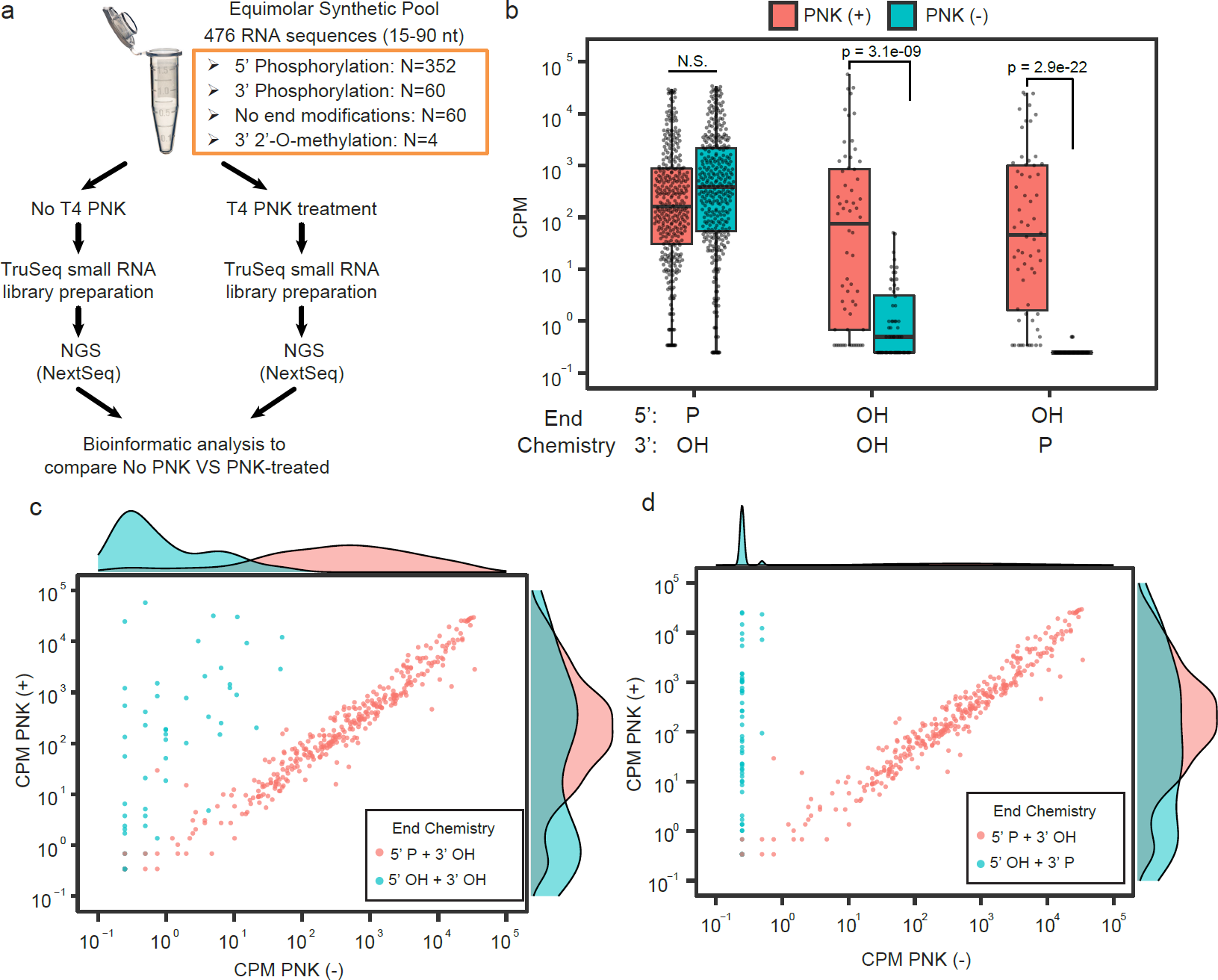
A modified protocol overcomes the low efficiency of standard small RNA library preparation methods for cloning short RNA sequences lacking 5’ phosphorylation**. 1A)** Schema of experimental design**. 1B)** Boxplots summarize the mean CPM observed for sequences contained in the synthetic equimolar pool sequences (*y-axis*, log_10_ scaled) presenting different end modifications (*x-axis*), as measured from libraries prepared using a standard ligation based small-RNA protocol and the phospho-sRNA-seq strategy. Boxes represent the mean +/-interquartile range (IQR), and whiskers represent 1st/3rd quartile 1.5 * IQR. Boxplots summarize mean CPM values for n = 352 (5’ phosphorylated + 3’ OH sequences), n= 60 (5’ OH + 3’ OH sequences) and n = 60 (5’ OH + 3’ phosphorylated sequences). Significant bonferroni-adjusted p-values are shown from one-tailed Wilcoxon Rank Sum tests for differences in abundance between PNK (+) and PNK (-), for sequences with the end chemistries shown (alternative hypothesis PNK + > PNK -). **1C and 1D)** Scatter plots showing the read distribution (CPM) observed for sequences of the synthetic equimolar pool with 5’ phosphorylation (red dots) and **1C)** without end modifications (teal dots) or **1D)** with 3’ phosphorylation (teal dots) as measured from libraries prepared using a standard ligation-based small-RNA protocol (*x-axis*) and the phospho-sRNA-seq strategy (*y-axis*). Marginal density plots are included as a summary of the data.

As shown in ***Figure 1B***, both strategies were able to recover the majority of sequences with 5’ phosphorylation and 3’ OH in our pool, most of which are human miRNAs (Median CPM PNK: 160.7; Untreated: 388.3). In contrast, the phospho-sRNA-seq approach recovered sequences that either lacked 5’ phosphorylation (Mann-Whitney one-tailed Bonferroni-adjusted p-value = 3.1e-09) or had 3’ phosphorylation (Mann-Whitney one-tailed Bonferroni-adjusted p-value = 2.9e-22), which were largely undetectable by the standard methodology (**Figures 1B - 1D**). We confirmed that these differences were not due to differences in sequencing depth, as the untreated library generated 4.1 million aligned reads, compared with only 2.9 million in the PNK-treated library. These results confirmed that standard ligation-based small-RNA protocols are poorly suited for capturing non-miRNA species lacking 5’ phosphorylation and, especially, those presenting a 3’ phosphorylation.

### Phospho-sRNA-seq combined with a high stringency bioinformatic pipeline enables reliable detection of mRNA/lncRNA fragments in human plasma

After validating the efficiency of our phospho-sRNA-seq strategy for capturing mRNA and lncRNA fragments with a variety of end modifications in a setting where the ground truth is known (i.e. a synthetic pool of RNA), we aimed to design and test a pipeline that could enable reliable evaluation of mRNA and lncRNA fragments in real plasma samples, where the exRNA composition is unknown and the risk of false positive calling is higher. To this end, we obtained platelet-poor plasma from five healthy control individuals (demographic features are shown in ***Appendix Table S2***), prepared triplicate libraries for each individual using both standard small RNA-seq methodology (TruSeq kit) and phospho-sRNA-seq and performed multiplexed sequencing on a HiSeq platform (***Figure 2A***).

**Figure 2.**
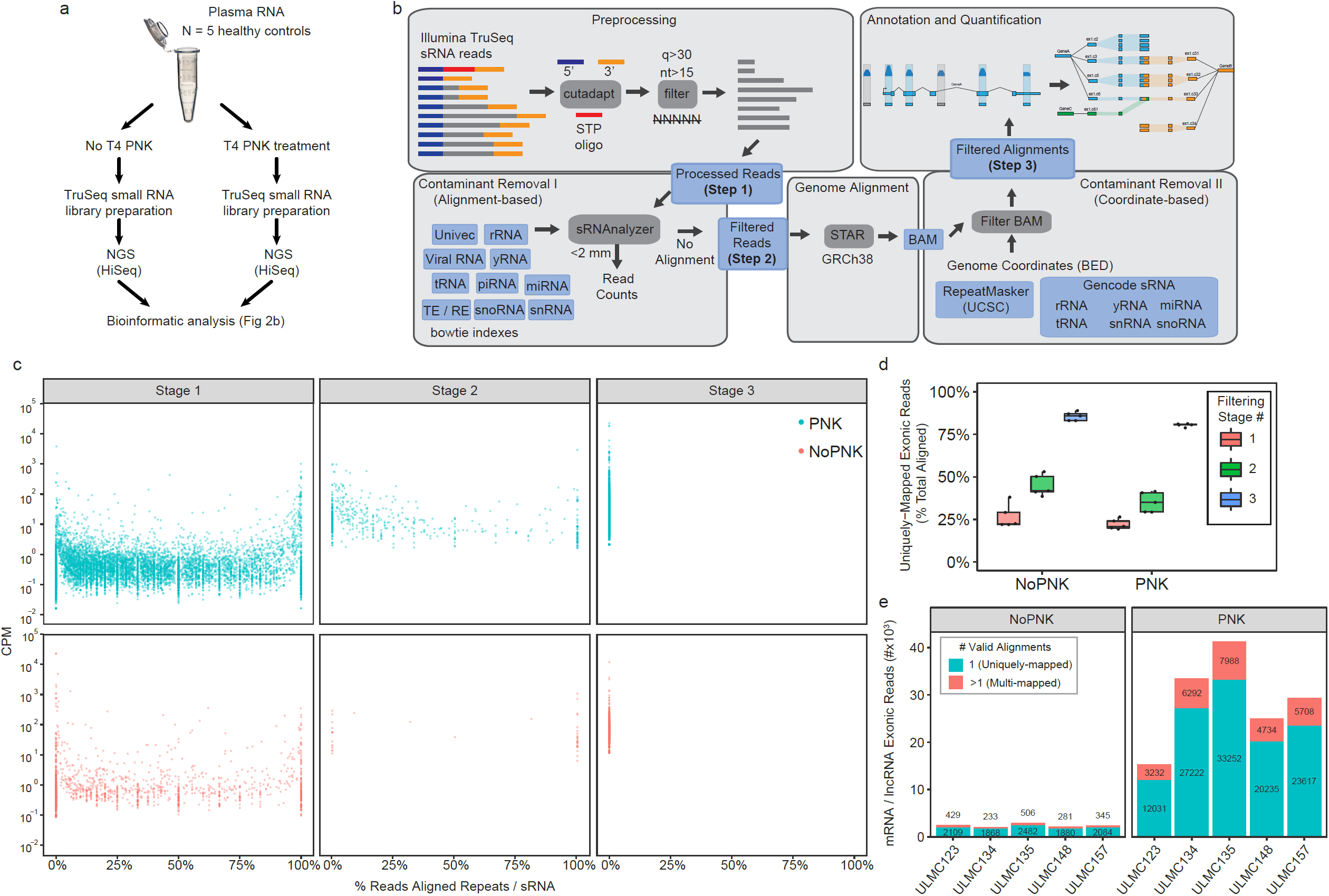
Phospho-sRNA-seq combined with stringent contaminant sequence filtering reduces false positive mRNA/lncRNA fragments. **2A)** Schema of experimental design. **2B)** Schema of bioinformatic analysis pipeline. **2C)** Scatter plot showing the percentage of reads aligned to repeats and small RNAs (*x-axis*) for the each filtering stage of our custom pipeline. Dots represent the mean CPM calculated for each gene across the five healthy control individuals. **2D)** Boxplots show the fraction of genome alignments that are unambiguously-aligned to mRNA and lncRNA exons, shown as the percent total reads aligned at each filtering stage. The points represent and the boxplots summarize the percentages calculated from combining alignments from three technical replicates for each of the (n = 5) healthy individuals. Boxes represent the mean +/-IQR, and whiskers represent 1st/3rd quartile 1.5 * IQR. **2E)** Barplots show the number of uniquely-mapped (teal) and multi-mapped (red) mRNA and lncRNA exonic reads remaining after the final filtering stage (Stage 3). Counts are plotted for each of the five healthy control individuals (*x-axis*) both in untreated (left panel) and PNK treated (right panel) samples. The count values are shown along with the corresponding bar plots.

Initial attempts to characterize mRNA and lncRNA fragments directly from adapter-trimmed and length-filtered reads revealed that non-mRNA/lncRNA sequences, including various endogenous small RNAs (rRNAs, tRNAs, miRNAs, etc) and repetitive elements, were leading to false positive detection and over-estimation of mRNA and lncRNA fragment abundances. To uncover relevant biological signal derived from mRNAs and lncRNAs, we developed a custom pipeline (**Figure 2B**) that employs multiple distinct filtering steps aimed at quantifying and removing potential sources of false signal, to enable the reliable detection of short mRNA and lncRNA fragments. Stage 1 of the pipeline involves trimming adapters, removing low-quality bases, and eliminating reads shorter than 15 nucleotides (see Materials and Methods for additional criteria). Next, the sRNAnalyzer pipeline(Wu *et al*, 2017) was adapted to quantify and remove reads aligning to any one of several sequence libraries containing exogenous RNAs (bacterial, fungal and viral), various “small endogenous RNA” sequences (defined here to include rRNAs, tRNAs, miRNAs, etc. as derived from Gencode and described in detail in Methods), and other possible contaminants (transposons, repetitive elements and Univec contaminants) (Stage 2). Reads with no valid alignments to these sequence libraries in Stage 2 are then aligned to the human genome. In Stage 3, genomic read alignments are filtered if found to have any overlap with RepeatMasker (UCSC) and small endogenous RNA annotation coordinates. This additional coordinate-based filtering step catches reads that were missed by the sRNAnalyzer workflow (see Materials and Methods for additional details).

As shown in ***Figure 2C***, without any filtering of small RNA and repeat-mapping reads (Stage 1), thousands of mRNA and lncRNA genes were falsely detected, or detected at artificially high levels due to a preponderance of reads aligning to transcript-embedded small endogenous RNA or repetitive element sequences. The pre-alignment to sequence databases in Stage 2 and the coordinate-based filtering in Stage 3 provided a step-wise removal of false positives from these endogenous sources (***Figure 2C***). Accordingly, the percentage of reads uniquely mapped to mRNA and lncRNA exons also increases through the sequential filtering stages of our pipeline (***Figure 2D***). Therefore, our analysis demonstrates the importance of stringent filtering of structural small RNA and other repetitive sequences, as failure to do so resulted in false-positive detection of many mRNA/lncRNA transcripts. It is also worth noting that sequences from libraries prepared with phospho-sRNA-seq mapped more frequently to mRNA and lncRNA exons, than those prepared using standard small RNA-seq, with the former showing a 10-fold increase of mRNA/lncRNA exonic reads on average (***Figure 2E).***

### Standard ligation based small RNA-seq pipelines are prone to false positive calling of mRNA/lncRNA fragments in human plasma

To evaluate how reliable standard small RNA-seq pipelines are for calling short mRNA and lncRNA fragments, we processed the sequencing data from healthy individual plasma exRNA through exceRpt, a pipeline specifically designed for the analysis of exRNA small RNA-seq data that uses its own alignment and quantification engine to map and quantify a range of RNAs including mRNA and lncRNA (see Materials and Methods). We then selected the 50 most abundant mRNA transcripts called by the exceRpt pipeline for evaluation through each stage of our custom high-stringency pipeline. As in our own pipeline, exceRpt first aligns adapter-trimmed reads to several small RNA databases for quantification, and only reads with no valid small RNA alignments are subsequently aligned to the human genome and used for mRNA and lncRNA quantification. However, tracking the reads corresponding to the top 50 most abundant exceRpt mRNA transcripts through our own high-stringency pipeline (***Figure 3A***) showed that although all 50 transcripts were detectable at Stage 1, most of them were filtered out or significantly reduced in relative expression, by subsequent filtering steps in Stages 2 and 3. We confirmed that a high proportion of them corresponded to small endogenous RNA species or repeat-mapping reads that were, therefore, ultimately filtered out in our pipeline (***Figure 3B***). Interestingly, for the libraries prepared using phospho-sRNA-seq, only 10 of the top 50 exceRpt transcripts were filtered out when analyzed through our highly stringent pipeline, as compared to 35 with standard small RNA-seq. These results demonstrate that standard small RNA-seq pipelines, even thoughtfully-designed ones like the exceRpt pipeline, which seek to map and remove some irrelevant small RNA and repetitive sequence species prior to alignment, are prone to false positive calling of mRNA and lncRNA fragments, thus limiting reliable identification of these exRNA species in plasma samples.

**Figure 3.**
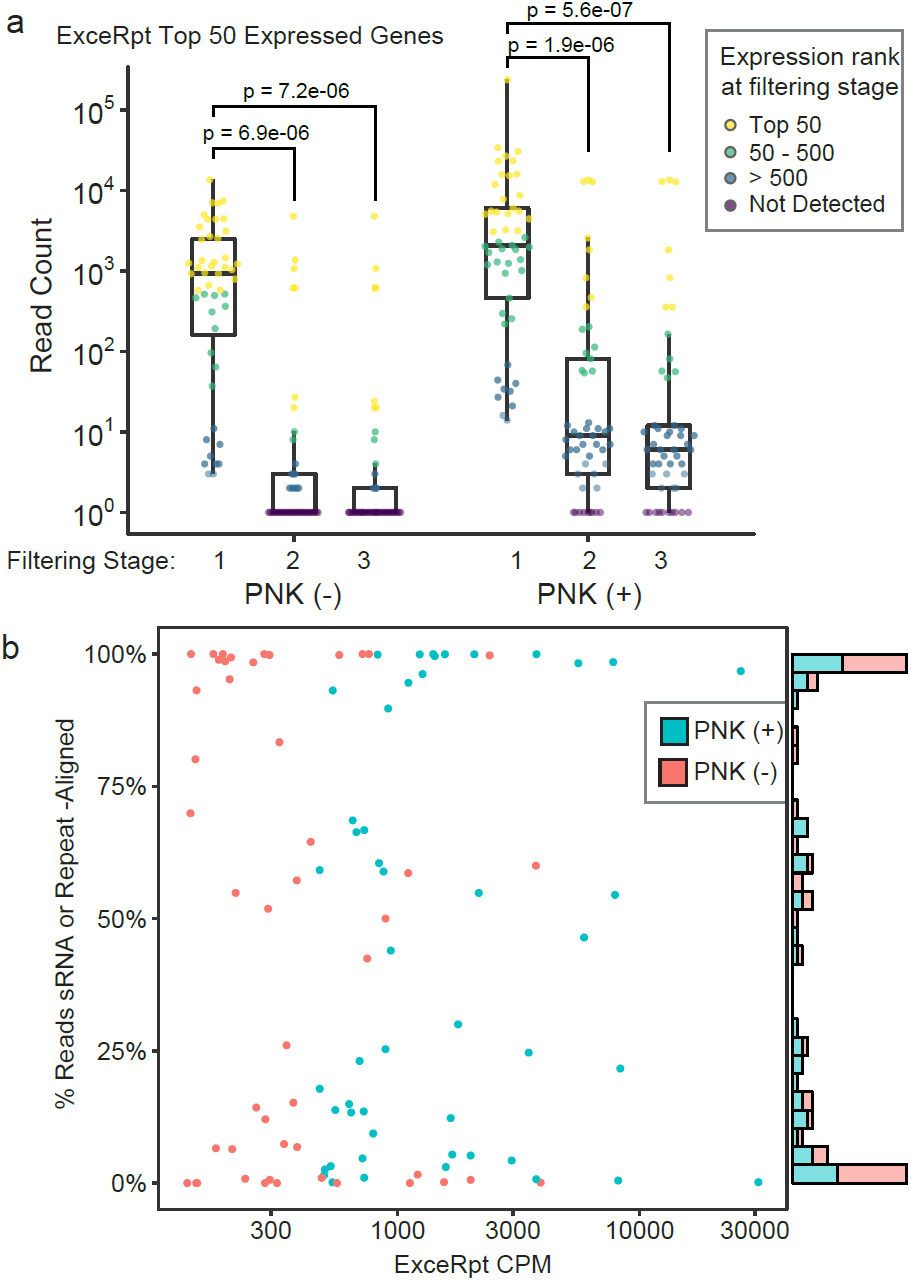
Re-evaluation of top transcripts called by a standard small RNA-seq analysis pipeline using our custom high stringency analysis pipeline. **3A)** The exceRpt exRNA-seq pipeline was used to analyze healthy subject plasma RNA (subject ULMC135) and the 50 most highly-expressed protein-coding mRNAs were quantified using our pipeline. Boxplots summarize the read counts measured when processed through our repeat filtering stages. Results from both PNK treated and untreated samples *(x-axis*) are shown. Gene abundances are shown as log_10_ read counts + 1 (*y-axis*). Individual points are color-coded by the rank of the gene expression observed at the stage indicated (rank 1 = highest expressed). Boxes represent the mean +/-IQR, and whiskers represent 1st/3rd quartile 1.5 * IQR. Bonferroni-adjusted p-values are shown from Wilcoxon Rank-Sum tests comparing the gene expression ranks in filtering stage 1 versus stage 2 or stage 3. **3B)** Scatterplot shows the CPM values reported by the exceRpt pipeline for the 50 most highly expressed mRNA or lncRNA genes (*x-axis*), versus the percentage that we found to overlap RepeatMasker or sRNA annotations (*y-axis*). Values are plotted for ULMC135 samples + PNK (teal) and - PNK (red).

### Assessment of short mRNA/lncRNA fragments in human plasma using phospho-sRNA-seq and our custom, high-stringency bioinformatic pipeline

After having validated that phospho-sRNA-seq combined with a custom, high-stringency bioinformatic pipeline enables reliable identification of short mRNA and lncRNA fragments in plasma samples, we sought to assess the abundance and features of these exRNA species in human plasma. In order to further substantiate the validity of these cell free mRNA fragments, we assessed the relative enrichment of reads aligning in the sense versus antisense orientation. The ligation-based library preparation we used ensures that that majority of reads are “stranded” -- that is, they should align in the same orientation as the transcript of origin. Contaminant-sequence alignments, or other noise introduced by sequencing artifacts is expected to be distributed more randomly, and would result in a more equal distribution of sense/antisense alignments. Thus, as a quality check, we confirmed that the exonic alignments of our plasma exRNA sequence reads were enriched for the sense orientation, relative to antisense, for mRNAs and lncRNAs (**Figure 4A**). The degree of enrichment for the sense orientation of lncRNAs was lower than for mRNAs, but this may be because the lncRNA database we used includes a diversity of lncRNA types (see Materials and Methods), including those overlapping mRNA transcripts on the opposite strand. As expected, sense strand bias was less evident for reads aligning to introns or promoter regions of mRNA or lncRNA genes (**Figure 4A**). We, therefore, focused our analysis of plasma exRNA on reads aligning to exons of mRNA and lncRNA genes.

**Figure 4.**
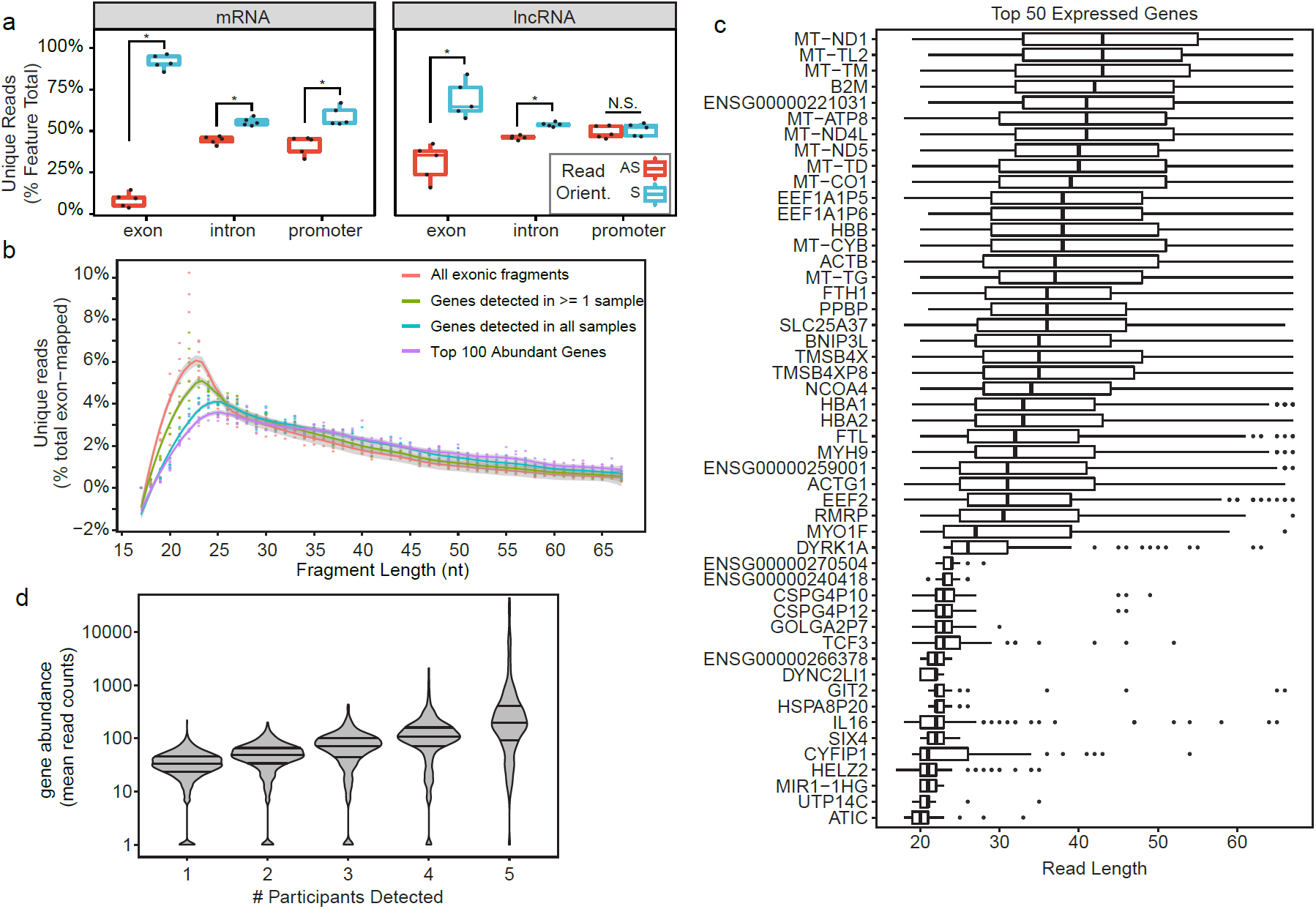
Assessment of short cell-free mRNA/lncRNA fragments in human plasma using optimized library preparation and analysis methods. **4A)** Boxplots showing the percentage of unambiguously annotated reads (*y-axis*) for mRNA and lncRNA exons, intron and promoters located in the sense and antisense strand as measured by phospho-sRNA-seq in plasma samples from healthy controls (N=5). **4B)** Read length distribution of exon-aligned reads in plasma samples from healthy controls (N=5) prepared with phospho-sRNA-seq. Read length is shown on the *x-axis*a nd the percent of exon-aligned reads shown on the *y-axis*. Dots represent percentages calculated for each of the five healthy control individuals. A smoothed trend line is shown and color coded based on the categories indicated. **4C)** Boxplots summarize the read length (*x-axis*) distributions for the 50 most highly-abundant genes (*y-axis*) across the five healthy control samples. Genes are sorted by median read length. **4D)** Violin plots showing gene abundance expressed as mean read counts (*y-axis*) as a function of the number of participants where they were detected (*x-axis*).

We found that our strategy is able to uncover thousands of mRNAs and lncRNAs present in physiological conditions in plasma from healthy individuals (N=5 healthy controls) (***Appendix Table S3)***. We evaluated the read distribution of the mRNA and lncRNA fragments identified in human plasma with our approach (***Figure 4B***) and found that on average they are fairly short (i.e. 20-25 nt range predominantly). However, it is worth mentioning that when we focused our analysis on the top 100 expressed mRNA and lncRNAs or on those mRNA and lncRNA expressed in all the samples, they tended to be slightly longer than the overall population of mRNA/lncRNA fragments, suggesting that longer read lengths are more frequently associated with more abundant and consistently-detected genes (***Figure 4B, 4D***).

Among the mRNA and lncRNA fragments we found in healthy individuals were these: (i) red blood cell-derived transcripts including several types of hemoglobin transcripts (e.g. HBA1, HBA2 and HBB); (ii) platelet-derived transcripts such as platelet-derived growth factors (e.g. PPBP); (iii) ubiquitous, highly expressed transcripts such as ferritin chains (i.e. FTH1 and FTL), mitoferrin-1 (i.e. SLC25A37), conventional non-muscle myosin (i.e. MYH9), multiple mitochondrial transcripts (e.g. MT-TL2, MT-ND1,MT-TM, MT-TD) and actin transcripts (e.g. ACTB and ACTG1) and; (iv) immune-related transcripts such as MHC class I molecules (e.g. B2M), interleukins (e.g. IL-6) and myosin IF (MYO1F), and (v) the lncRNAs MALAT-1 and NEAT1 (***Figure 4D and Appendix Table S3***). As expected, the mRNA and lncRNA fragments that were the most consistently detected across multiple individuals were also the most highly abundant ones (***Figure 4D)***.

### Pathophysiologic processes are reflected in plasma exRNA transcriptome profiles revealed by phospho-sRNA-seq

Having confirmed that the phospho-sRNA-seq approach with high-stringency bioinformatic analysis enables the detection of mRNA and lncRNA fragments consistently expressed in plasma from healthy individuals under physiological conditions, we next sought to evaluate if pathophysiological processes could be detected as changes in tissue-specific or enriched mRNA and/or lncRNA fragment exRNAs. To this end, we collected serial plasma samples from patients undergoing allogeneic bone marrow transplantation (BMT) (N=26 samples from 2 different patients), prepared phospho-sRNA libraries from each time point and performed multiplex sequencing using a HiSeq platform (***Figure 5A***).

**Figure 5.**
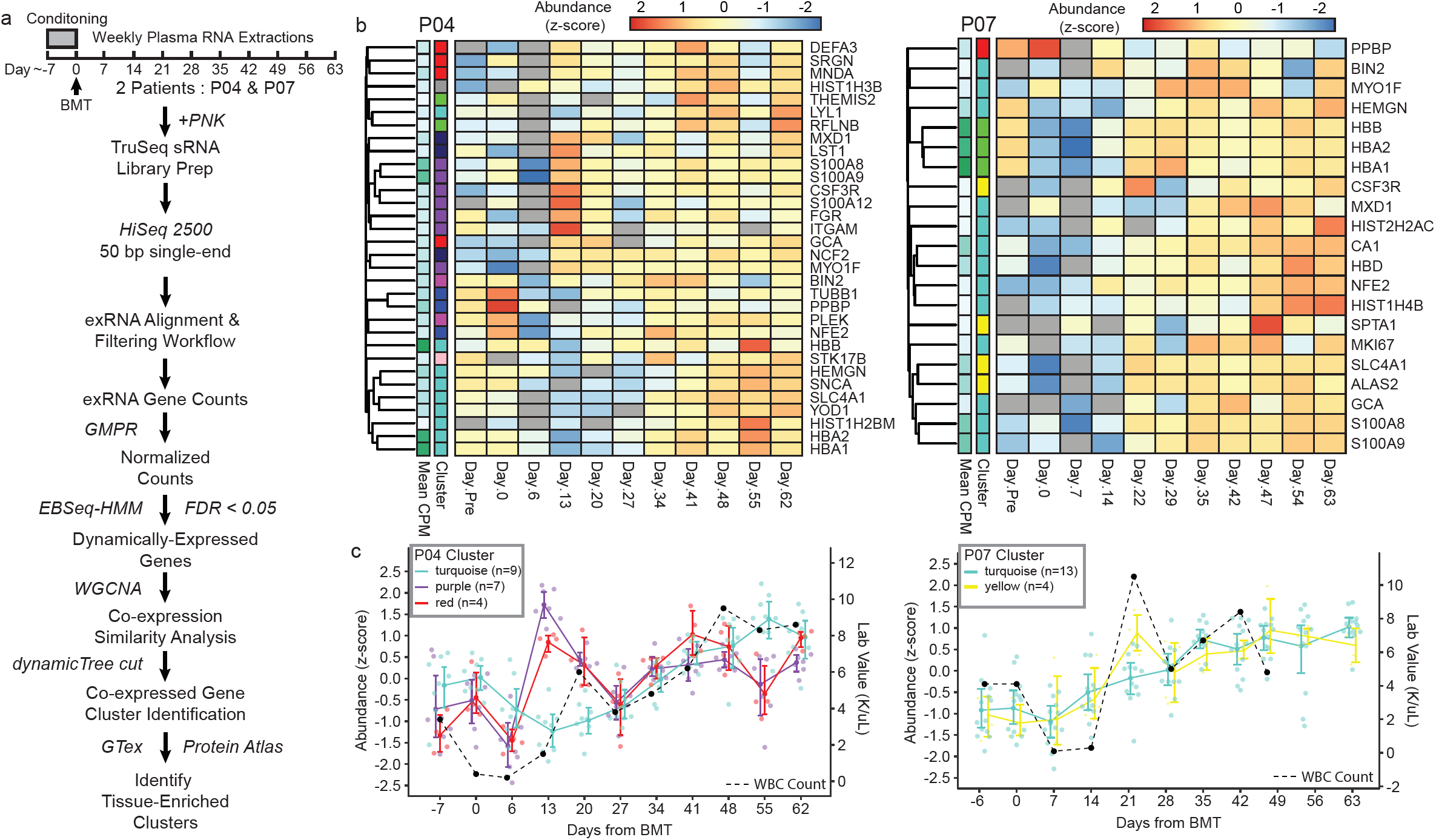
Cell-free mRNA/lncRNA fragments inform about pathophysiological processes in patients undergoing allogeneic bone marrow transplantation. **5A)** Schema depicts timing of the serial sample collection, experimental methodology and bioinformatics and enrichment analysis. **5B)** Heatmaps show the expression patterns of bone marrow-enriched genes that were detected in patients P04 (left) and P07 (right) datasets, and found differentially abundant by EBSeq-HMM (FDR < 0.01). GMPR-normalized read counts were centered gene-wise to have a mean of 0 and standard deviation of 1. The “cluster” row annotations indicate the co-expression cluster identified by WGCNA. P04 clusters turquoise, purple and red, and P07 clusters turquoise and yellow were significantly enriched for bone marrow transcripts (hypergeometric test; FDR < 0.01). **5C)** Graphs show the expression patterns for the co-expression clusters significantly enriched for bone marrow-enriched transcripts. Individual points are shown for each gene in the P04 (left) and P07 (right) co-expression clusters, and are colored according to the cluster IDs. Colored lines indicate the mean expression of each cluster, and error bars represent a bootstrapped (B=1000) 98% CI of the mean. Black dashed lines indicate the white blood cell counts obtained from lab results measured on the same day.

For the sequence data analysis (***Figure 5A***), we reasoned that in order for mRNA/lncRNA exRNA sequences in plasma to have potential as biomarkers, we would see patterns in specific sets of genes that correlate over time to biological processes happening within the patients. We began our analysis by using the EBSeq-HMM R package, which uses an autoregressive Hidden Markov modeling strategy to test for genes that show evidence of differential expression over time, using an FDR cutoff of 0.01. We hypothesized that the differentially-expressed genes with similar expression patterns have similar tissue origins or biological functions. We found sets of differentially expressed genes showing concordant temporal patterns (***Appendix Table S4***), some that were significantly enriched for bone marrow-specific transcripts (***Figure 5B and Appendix Table S4***) and others for liver-enriched transcripts (***Figure 6A and Appendix Table S4***). Since bone marrow transplantation is a process that can be followed through the peripheral white blood cell count (WBC), we plotted abundance of bone marrow-specific exRNA transcript fragments over time along with WBC count. As shown in ***Figure 5C***, we saw that transcript fragments corresponding to the bone marrow gene set tracked with bone marrow reconstitution, initially declining as expected during the period of early neutropenia in the first week after transplant followed by a rise corresponding to recovery of the WBC count. We wondered whether the liver-enriched transcripts that varied over time (i.e., those identified above) would track temporally with liver injury, which is common in BMT patients, sometimes as a side effect of medications given as part of their clinical care. By plotting blood levels of serum aminotransferases (AST and ALT), two proteins produced by liver cells that are used clinically for detecting liver injury, together with levels of exRNA liver-specific transcript fragments over time, we saw that levels of the liver-specific RNA fragments generally correlated with changes in AST/ALT (***Figure 6B***). Thus, we concluded that bone marrow-specific and liver-specific exRNA transcript fragments show distinct expression patterns corresponding to known biology and relevant, established clinical laboratory markers. These results provide a proof-of-concept that this approach can provide access to a circulating transcriptome with potential for biomarker development.

**Figure 6.**
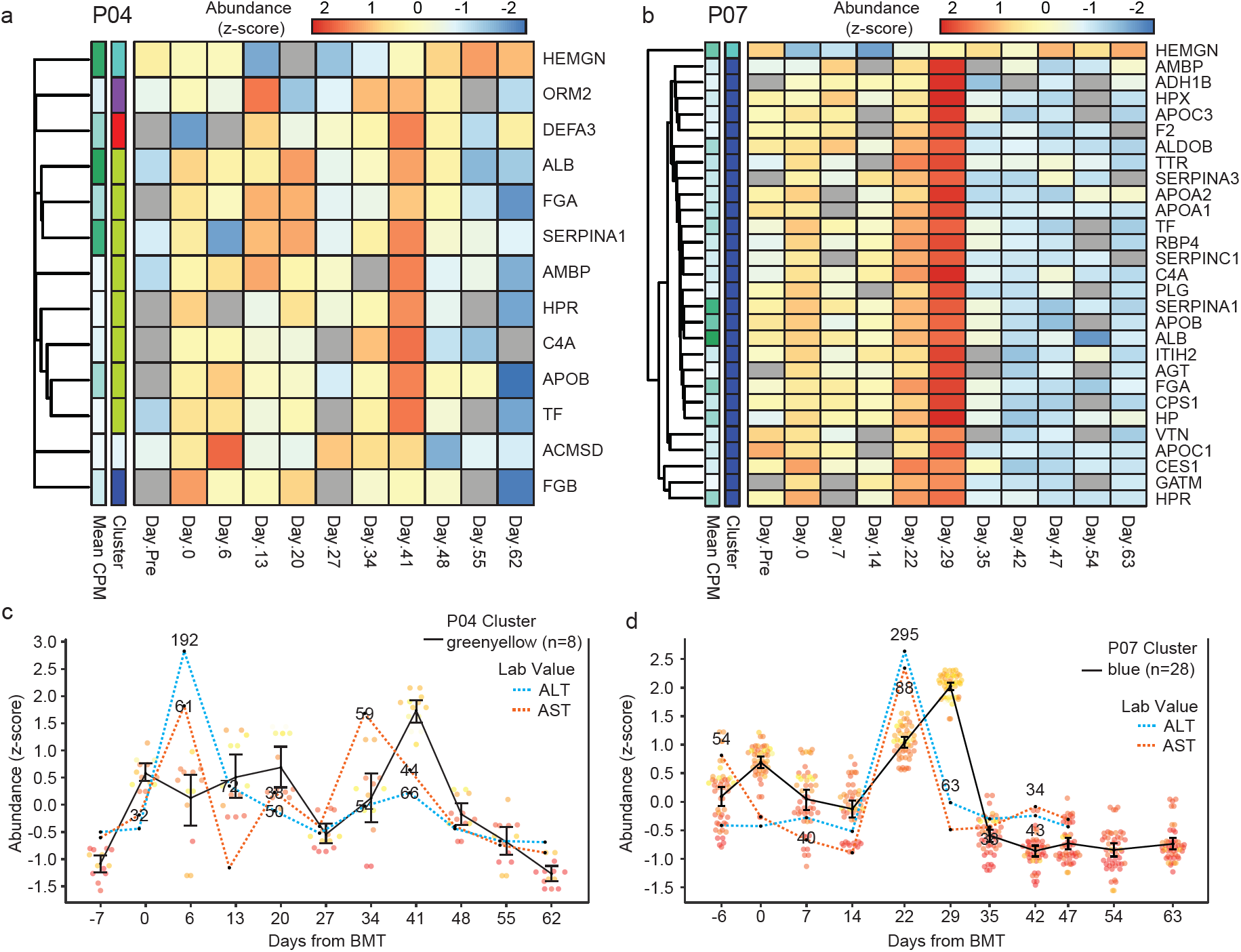
6A) Heatmap of liver-enriched genes detected and found differentially-abundant by EBSeq-HMM (FDR<0.01) in **6A)** P04 and **6B)** P07 samples. The “cluster” row annotations indicate the co-expression clusters identified by WGCNA. P04 cluster, greenyellow, and P07 cluster, blue, were significantly enriched for liver-specific and enriched transcripts (hypergometric test; FDR < 0.01). **6C)** Graph shows the expression for genes in the P04 co-expression cluster, greenyellow, along with AST and ALT lab values taken on the same day. **6D)** Graph shows the expression of P07 co-expression cluster, blue, along with AST and ALT lab values. Points represent the z-transformed log_2_ read counts for each of the genes in the cluster (P04 greenyellow: n=8; P07 blue: n=28). The black lines indicate the mean expression of each cluster and error bars represent a bootstrapped (B=1000) 98% CI of the means. Blue and red dashed lines represent lab values for AST and ALT liver enzymes, respectively. AST and ALT levels were centered to a mean of 0 and standard deviation of 1 for plotting. The actual lab values are shown for all elevated readings (AST > 30; ALT > 35).

## Discussion

There has been rapidly growing interest in the study of exRNAs, both for their potential clinical application as biomarkers of disease measurable in biofluids such as blood(Roser *et al*, 2018; Schwarzenbach *et al*, 2011), as well as for their potential biological functions(Zhang *et al*, 2010; Shah & Calin, 2014; Hu *et al*, 2012). Most studies of exRNAs in human biofluids to date have focused on miRNAs. Unlike miRNAs, which are a small fraction of total genes, lncRNAs and mRNAs comprise a majority of the transcriptome and hold great potential as biomarkers given their exquisite tissue specific expression(Iyer *et al*, 2015; Liu *et al*, 2008b) and the fact that mRNA gene signatures in tissues have proven to be powerful biomarkers in different clinical settings(Perou *et al*, 2000; Potti *et al*, 2006; Ben-Porath *et al*, 2008; Chen *et al*, 2007). Thus, the ability to read transcriptomic information through exRNA profiles in blood plasma is important for enabling for clinical applications.

However, mRNAs and lncRNAs have not been easily or consistently detectable as exRNA in blood plasma. RNAs found in plasma have generally been seen to be fragmented relative to their cellular RNA counterparts(Mitchell et al. 2008), and most studies have used small RNA-seq protocols that are designed to sequence miRNAs. These protocols typically employ RNA ligase-based adapter ligations, followed by reverse transcription and PCR to generate libraries for high throughput sequencing. Such protocols commonly rely on the presence of 5’-phosphate and 3’-hydroxyl ends on the target RNA(Hafner *et al*, 2008), which are produced by the RNase III class of ribonucleases, including the double-stranded ribonuclease Dicer that is responsible for processing precursor miRNAs to generate mature miRNAs(Knight & Bass, 2001; Ha & Kim, 2014). We hypothesized that mRNA and lncRNA transcripts in the blood circulation are likely to be acted upon by different classes of ribonucleases that may not produce ends conducive to sequencing by standard small RNA-seq library protocols. Specifically, abundant RNases in human circulation such as those belonging to the ribonuclease superfamily A(Lu *et al*, 2018) degrade RNA dinucleotide bonds leaving a 5′ hydroxyl and 3′ phosphorylated product (Cuchillo *et al*, 2011), thus rendering cleavage products unsuitable for standard ligation-based library preparation protocols.

We sought to test this hypothesis using both “ground truth” samples of synthetic RNA pools with variable 5’ and 3’ end modification states and biological (plasma) samples. Our results clearly showed that T4 PNK treatment vastly increased the recovery of fragments either lacking a 5’-phosphate or having a 3’-phosphate. This is consistent with the properties of T4 PNK, which has both 5’ kinase and 3’ phosphatase activities(Cameron & Uhlenbeck, 1977; Richardson, 1965; Novogrodsky & Hurwitz, 1966). Moreover, these results highlight that standard ligation-based small RNA-seq approaches are not well suited to characterize exRNA beyond miRNAs and other sequences sharing the same end chemistries. Our method revealed that human plasma contains abundant mRNA and lncRNA transcript fragments, corresponding to thousands of human genes. Thus, mRNA and lncRNA fragments are a substantial component of the extracellular transcriptome in human plasma. Our incorporation of synthetic RNA pools as a ground truth reference was especially important in our study, as it allowed us to demonstrate clearly that lack of 5’ phosphorylation and presence of 3’ phosphate are both impediments to recovery of these fragments by standard small RNA-seq protocols.

A key lesson learned from our study is the importance of a highly stringent data analysis for accurate identification of mRNA and lncRNA fragments from phospho-sRNA-seq sequence data. This is because these relatively short sequences frequently align to multiple locations in the genome and RNAs arising from repetitive DNA transcription or non-coding small RNAs such as ribosomal RNA fragments, can spuriously align to mRNA and lncRNA exons. To avoid these false positive calls, we developed a three step filtering pipeline specifically designed for the reliable identification of mRNA and lncRNA fragments. To date, pipelines for the analysis of small RNA-seq data such as the exceRpt pipeline (http://genboree.org/theCommons/projects/exrna-tools-may2014/wiki/Small_RNA-seq_Pipeline) have been designed with a predominant focus on miRNA annotation. Perhaps not surprisingly, we found that when applied to phospho-sRNA-seq data analysis for mRNA/lncRNA identification, there was a high rate of false positive annotation. It is possible that this phenomenon may have affected prior results of plasma exRNA profiling using non-PNK-based approaches which although focused primarily on miRNAs, also commented on the finding of mRNA and lncRNA transcripts in plasma(Max *et al*, 2018; Yuan *et al*, 2016; Huang *et al*, 2013). Future work could determine the extent to which reports of mRNA and lncRNA fragments from libraries not incorporating PNK may reflect false positive mis-annotation, due to limitations of bioinformatic analysis pipelines. We propose that the high-stringency pipeline we describe here is one approach for mitigating the false positive rate, and there may be further bioinformatic approaches that can also be developed.

We must acknowledge, however, that the strict filtering used to remove repeats, although effective in reducing false positives, also potentially removes some valuable information and can lead to false negatives. Future work could utilize additional features such as high-confidence alignments throughout the exons of a gene in the proper orientation relative to the transcript. This may provide enough evidence to enable us to capture reads aligning to embedded repetitive sequences in bona fide mRNA/lncRNA transcripts, for example.

T4 PNK is a well-characterized enzyme frequently used to create appropriate and homogeneous RNA ends before long RNA sequencing. Specifically, this enzyme has been used in long RNA sequencing protocols to render tissue RNA suitable for adapter ligation after experimental heat- or alkali-based fragmentation(Lee *et al*, 2013; Lamm *et al*, 2011). T4 PNK has also been included in protocols aimed to identify 5’PPP moieties that correspond to transcriptional start sites in bacteria (i.e. differential RNA-seq)(Vvedenskaya *et al*, 2015). In contrast, T4 PNK is not generally used as part of small RNA protocols and it has not been specifically evaluated as an strategy for revealing novel mRNA/lncRNA fragment sequences in biological samples such as plasma, which are missed by standard ligation-based small RNA protocols.

We applied phospho-sRNA-seq to analyze longitudinal plasma samples from bone marrow transplant patients for two reasons. The first was to provide additional validation for our methods and data analytic pipeline, in that finding gene signatures in plasma that correspond to known time-dependent biological changes in patients would make it highly unlikely that the mRNA/lncRNA fragments we found were spurious and due to mis-annotation, for example. The second was to establish the proof-of-concept that mRNA/lncRNA fragment gene signatures in plasma correlate with human biology, indicating the potential of this method as a new, broadly applicable “liquid biopsy” approach. As shown here, we found through an analysis beginning with temporal plasma exRNA gene expression profiles and progressing to identify tissue-specific gene sets over-represented in the dynamic profiles, that specific gene signatures corresponding to bone marrow and liver both tracked in a manner that paralleled the dynamic processes of bone marrow reconstitution and liver toxicity, respectively. These results indicate that the phospho-sRNA-seq approach, when combined with high-stringency bioinformatic data analysis, could be applied to a range of diseases in which tissue injury figures prominently (e.g., drug toxicities, immune-mediated damage such as in autoimmune conditions or side effects of cancer immunotherapy, etc.). This is likely true of any process or pathology that releases transcriptome elements into the blood circulation (e.g., cell death occurring in growing tumor tissue).

Although most prior plasma exRNA studies have focused on miRNAs, there are some studies, using methods other than phospho-sRNA-seq, that have reported mRNAs in the plasma. One notable report using a random primer cDNA synthesis long RNA-seq protocol, detected mRNAs and lncRNAs in plasma(Koh *et al*, 2014). Given that the minimal length of RNAs general captured by that protocol is larger than what we observed with phospho-sRNA-seq, we expect that approach is unlikely to have captured many of the mRNA/lncRNA fragments found in our study. Taken together, the results suggest that exRNA transcripts of a broad range of lengths might coexist in the human circulation. It remains an open question at this point as to what fraction of the plasma exRNA transcriptome is present as shorter fragments of the form revealed in our study, or as the presumably longer RNAs reported in prior studies. The shape of the exRNA size distribution observed in our data would indicate that longer fragments (e.g., > 100 bases) may exist, but may be a minority. However this question would need to be addressed in future studies to obtain a more accurate characterization of the exRNA transcriptome present in blood and other biofluids.

Although our phospho-sRNA-seq strategy allowed us to expand the spectrum of sequences that can be detected in plasma (i.e. sequences lacking 5’ phosphorylation and/or presenting a 3’ phosphate end), there are certainly still limitations of this approach. Even though we expect that most of the cleavage products derived from mRNA/lncRNA in circulation will present 5’ OH and 3’P due to the abundance of superfamily A RNAses in plasma, it is possible that there are exRNAs in plasma with end groups that are not addressed by our methodology (e.g. 5’ cap). Second, it is worth mentioning that the efficiency of phospho-sRNA-seq for identifying mRNA/lncRNA fragments is affected by the recovery of other abundant plasma fragments such as ribosomal RNA and Y RNA that can dominate the sequencing data, thus reducing the depth of sequencing available for detecting mRNA and lncRNA fragments. To this end, we foresee that in the future, the sensitivity for detecting relevant mRNA and lncRNA fragments in plasma might be improved by designing specific methods for depleting abundant, undesired fragments such as fragments of rRNAs and Y RNAs, or for enriching for fragments corresponding to panels of selected transcripts. It is clear that there is still room for method improvement and there may be many more mRNA and lncRNA fragments in plasma than we have identified here. It is worth noting that new ligation-free strategies have been developed recently for small RNA-seq (Turchinovich *et al*, 2014). Future studies will be required to determine the efficiency and effectiveness of these approaches for detecting mRNA and lncRNA fragments in circulation. Moreover, taking into account our results, we envision that sequence data analytic pipelines specifically designed for reliable analysis of mRNA/lncRNA fragments, such as the one described here, will be required for the analysis of exRNA sequencing data generated with these strategies.

In summary, our results highlight that there is greater complexity of the extracellular transcriptome in human biofluids than previously known and that phospho-small RNA-seq can provide access to transcriptomic signatures in plasma that are inaccessible by standard small RNA-seq methods. The methodology presented here provides access to a new class of extracellular RNAs for development as liquid biopsy biomarkers for a variety of diseases. In addition, it may be useful to investigate this technique for application to other settings where RNA is highly fragmented, including formalin-fixed paraffin-embedded archival tissue specimens, as well as extremely old specimens of cells or tissues, where RNA may likewise be present in highly fragmented form.

## Materials and Methods

### Synthetic reference sample

A synthetic equimolar pool containing 476 synthetic RNA oligonucleotides was prepared in an RNase-free environment and working on ice to minimize degradation. The pool was prepared by combining (i) 286 human miRNAs (ii) a set of 190 additional, custom-synthesized RNA oligonucleotides, to generate the pool in which each of the 476 RNA oligonucleotides is present at equimolar concentration. The latter set of 190 comprises miRNAs and non-miRNA sequences of varied length from 15 to 90 nt, which were synthesized, HPLC purified and quantified spectrophotometrically by IDT. The pool of RNA oligonucleotides is available to qualified investigators seeking to reproduce the synthetic equimolar for non-commercial purposes, by request of the corresponding authors (as long as supplies last). The resulting equimolar pool was aliquoted in prelabeled DNA-, DNase-, RNase-, and pyrogen-free screw cap tubes with low adhesion surface and stored immediately at -80 C. The complete list of RNA sequences comprising the equimolar pool is provided in ***Appendix Table S1***.

### Biological samples

Plasma samples from 5 healthy donors and serial plasma sample from 2 patients undergoing allogeneic bone marrow transplantation were collected in 10-mL K2EDTA plasma tubes (Vacutainer 366643; Becton Dickinson) and processed within one hour of blood draw following a two centrifugation protocol to obtain platelet-poor plasma as previously described(Cheng *et al*, 2013): (i) 3,400 xg at room temperature for 10 minutes with high brake; and (ii) 1940 xg at room temperature for 10 minutes without brake. Plasma was stored at -80C until RNA isolation. The University of Michigan IRB approved the study protocol to consent participants and collect samples. Informed consent was obtained from all subjects, and the samples were subsequently de-identified before distributing to the laboratory personnel generating the libraries. The studies conformed to the principles set out in the WMA Declaration of Helsinki and the Department of Health and Human Services Belmont Report.

RNA was isolated from 200 ul of plasma using the miRNeasy mini kit (Qiagen, Hilden, Germany) according to the manufacturer’s protocol with the following modifications. Plasma samples were mixed with five sample volumes of QIAzol reagent and vortexed for 10 s. Samples in QIAzol were incubated at room temperature for 5 min to inactivate RNases. Next 0.2 volumes of chloroform were added to each sample. At that point, the manufacturer’s protocol was followed.

### Library preparation and sequencing

The input for library preparation was 10 femtomoles of RNA for the synthetic equimolar pool and 5 μl of RNA for the biological plasma samples. Standard ligation-based small RNA libraries were prepared using the TruSeq small RNA kit (Illumina, San Diego, CA, US) according to the manufacturer’s instructions. Size selection was performed using pre-cast 6% acrylamide gels (Invitrogen, Carlsbad, CA, US) including all products from 140-200 bp plus any additional visible bands of greater size. To perform phospho-sRNA-seq, synthetic and plasma RNA samples were pretreated with T4 polynucleotide kinase (NEB, Ipswich, MA, US) using an RNA input of 7 ul in a final reaction volume of 10 ul and following the manufacturer’s instructions. After the enzymatic treatment, biological RNA samples were purified by performing sequential washes in silica columns (Zymo, Irvine, CA, US): (i) 900 ul of buffer RWT (Qiagen, Hilden, Germany); (ii) 900 ul of buffer RPE (Qiagen, Hilden, Germany); (iii) 900 ul of ethanol 200 proof, molecular biology grade (Fisher Scientific, Waltham, MA, US) and; (iv) 900 ul of 80% ethanol. Libraries were then prepared using the TruSeq small RNA kit according to the manufacturer’s instructions. Size selection was performed as described above. For the libraries generated from patients undergoing bone marrow transplantation we narrowed the range of size selection to 140-165 bp to reduce the abundance of contaminants such as Y RNAs.

Libraries were multiplexed and sequenced using the Illumina NextSeq 500 (synthetic equimolar pool) and Illumina HiSeq 2500 (healthy controls and patients undergoing allogeneic bone marrow transplantation) specifying 75 bp and 50 bp single-end runs, respectively.

## Computational Methods

### Synthetic Pool Library Analysis

Illumina NextSeq reads from equimolar synthetic pool libraries were processed to trim adapters, remove low-quality bases and filter short reads. Reads as short as 14 nt were allowed for detection of the shortest oligos in the pool. The sRNAnalyzer workflow was used in “multi” mode to align the processed reads to Univec contaminants, and a bowtie sequence database containing the 476 Equimolar Pool sequences. Read alignments were loaded into R for processing and analysis. Read counts for each sequence in the pool were based on reads with <= 2 mismatches, aligning in the sense (+) orientation. For reads with valid alignments to both Univec and Equimolar Pool sequence libraries, the alignment(s) with the fewest mismatches were used.

### exRNA Processing Pipeline

TruSeq adapters and stop oligo sequences were trimmed with cutadapt (v 1.91) using processing steps adapted from the sRNAnalyzer workflow(Martin, 2011; Wu *et al*, 2017). The sRNAnalyzer framework was also adapted to align adapter-trimmed reads 15 nt and longer to several sequence databases containing known small RNA families and contaminant sequences(Wu *et al*, 2017). A table with descriptions of the included sequences databases are provided in ***Appendix Table S5***. Up to two mismatches were allowed in the alignment. Reads that had no valid alignments to the small endogenous RNA and contaminant databases were aligned to the human genome (GRCh38) using STAR (v 2.5.0A)(Dobin *et al*, 2013). The following parameters were altered from default:

outFilterMultimapNmax=1000000; outFilterMismatchNoverLmax= 0.1; outFilterMatchNmin=15; outFilterMatchNminOverLread=0.9; outMultimapperOrder=Random; outSAMtype=BAM

Unsorted; outReadsUnmapped=Fastx ; outSAMattributes=All ;

outSAMprimaryFlag=AllBestScore; alignIntronMax= 1; alignIntronMin= 2; alignSJDBoverhangMin=999

These parameters remove the splicing-aware alignment capability and limit the extent of “soft-clipping” at the ends of the alignment.

### Multi-mapping scaling and gene quantification

Read counts were weighted using a strategy similar to that employed by CSEM, which gathers mapping information from neighboring read alignments to weight read counts towards loci with the most unambiguous mapping information(Chung *et al*, 2011). Our strategy differs in that we 1) gather mapping information from neighboring read alignments across all samples in the cohort, 2) restrict our search to directly overlapping fragments, and 3) retain the mapping ambiguity information to allow identification of commonly co-mapping genes. To do this, a bipartite network was created using the R package, igraph, to connect reads with all overlapping clusters of mapped loci (Csardi G, Nepusz T: The igraph software package for complex network research, InterJournal, Complex Systems 1695. 2006). All connected components were identified, using the mapping ambiguity information from all connected reads to weight reads more strongly to those regions with more unambiguously-mapped reads. Read alignments were annotated for overlap with A) Gencode transcripts, B) Gencode endogenous small, non-coding RNAs and C) RepeatMasker annotation coordinates (UCSC genome browser). All alignments were removed for any read aligning to Gencode small endogenous RNA or RepeatMasker loci (minimum 1 bp overlap in either orientation).

### Comparison with exceRpt pipeline

The exceRpt small exRNA analysis pipeline (v 4.6.2) implemented on the Genboree Workbench (http://genboree.org/java-bin/login.jsp) was used to process and analyze healthy control samples for, ULMC 135, prepared both using the standard TruSeq library protocol, and the modified phospho-sRNA-seq method.. Default exceRpt pipeline parameters were used, except to set the minimum read length to 16, and to specify TruSeq small RNA adapters for trimming. Post-processed files with Gencode read counts and reads per million were compared with the ULMC135 reads were collected (Stage 1) after adapter trimming and size filtering, (Stage 2) after sRNAnalyzer alignment and contaminant removal and (Stage 3) after genome alignment and removal of RepeatMasker and small endogenous RNA annotations. Reads from each stage were aligned to the human genome with STAR, using the same alignment parameters described above. These parameters were largely copied from those used by the exceRpt pipeline to make the alignments as comparable as possible. Comparison was limited to genes detected by both pipelines. Because the exceRpt pipeline output included only summarized gene abundance, the percent of small endogenous RNA or repeat-aligned reads were based on the alignments from our pipeline. An alignment was considered “sRNA or Repeat-Aligned” if any alignments for that read overlapped RepeatMasker or Gencode small endogenous RNA coordinates (minimum 1 nt overlap on either strand). Fragments were summarized at the gene level using muti-mapping-weighted exon-aligned fragments, comparable to that used by the exceRpt pipeline for gene-level quantification.

### Cell free RNA enrichment in coding and non-coding regions of mRNA and lncRNA transcripts

Exon, intron and promoter (2 kb upstream + 0.2 kb downstream nt) coordinates were extracted from Gencode (v27) protein-coding and long non-coding RNA annotations. RepeatMasker and sRNA-filtered genomic alignments from the five healthy individuals were intersected with these coordinates, requiring a minimum of bp of overlap in either orientation. Ambiguous annotations were allowed, but counted only once per unique combination of read and feature. Read alignments were considered “Sense” or “Antisense” based on the relative orientation of the read alignment and the mRNA or lncRNA feature. Read length distributions and gene abundance was calculated based on sense-aligned exonic reads.

### Analysis of Bone Marrow Transplant Cohort

HiSeq reads from P04 and P07 bone marrow transplant patients were processed and filtered as described above. Gene-level counts were calculated separately for P07 and P04 samples, using the multi-mapping-weighted read counts from mRNA and lncRNA exon-aligned read fragments. The resulting gene count matrices were normalized across samples using a robust Geometric Mean of Pairwise Ratios (GMPR) method, suitable for sparse data sets(Chen *et al*, 2018). The GPMR-calculated size factors were provided as input to the R package, EBSeq-HMM, which employs an autoregressive Hidden Markov Modeling strategy to identify genes with non-static expression dynamics over the course of the time series. EBSeq-HMM was run separately for P04 and P07 samples. An initial run was performed using a low number of iterations (n=5) to test a range of fold-change estimates (1.0 to 2.0, by 0.2). The estimate that maximizes the log likelihood was then used for a second run of the algorithm with a higher number of iterations (100). Significantly altered genes were selected at an FDR cutoff of 0.01(Leng *et al*, 2015). GMPR-normalized read counts from the significantly-altered genes were clustered using the WGCNA workflow(Langfelder & Horvath, 2008)

## Tissue Enrichment

Databases of tissue-enriched genes were obtained from GTex and Human Protein Atlas data curated by the TissueEnrich R package(Jain & Tuteja, 2018). Significant enrichment was determined with a hypergeometric test, and using a background of all genes used as input to EBSeq-HMM analysis.

## Acknowledgments

We thank T. Churay for assistance in obtaining clinical specimens, X. Cao and A. Chinnaiyan for assistance with sequencing of healthy control libraries, E. Sandford for assistance in specimen management, J. Vandesompele, E. Fearon, K.E.A. Max, T. Tuschl and B. Knudsen for helpful discussions, J.S. Rozowsky, R. Kitchen, S.L. Subramanian, W. Thistlethwaite and A. Milosavljevic for facilitating access to the exceRpt pipeline, C. Gates for assistance with the bioinformatics pipeline, and K. Wang for providing synthetic miRNA oligonucleotides. We acknowledge funding support from the NIH Extracellular RNA Communication Common Fund grants: U01 grants HL126499 to M.Tewari and HL126496 to D.J.G., and from the A. Alfred Taubman Medical Research Institute (Grand Challenge Award) to M.Tewari and S.W.C.. Research reported in this publication was also supported by the National Cancer Institute of the NIH under Award Number P30CA046592 by the use of the following Cancer Center Shared Resource at the University of Michigan: DNA Sequencing. M.D.G. acknowledges support from a Precision Health Scholar Award from the University of Michigan Precision Health Center and a Juan Rodes contract (JR18/00026) funded by the Spanish Institute of Health Carlos III from the Ministry of Economy and Competitiveness (co-funded by European Social Fund (ESF)). D.J.G. also acknowledges a special technology support award from the Pacific Northwest Research Institute to his lab. The content is solely the responsibility of the authors and does not necessarily represent the official views of the National Institutes of Health.

## Author Contributions

M.D.G. and M. Tewari conceived of the project. M.D.G. and A.E. designed and performed experimental work. M.D.G., R.M.S., A.E., S.W.C, D.J.G. and M.Tewari interpreted results. M.D.G., R.M.S., and M.Tewari wrote the manuscript. R.M.S. designed and implemented the computational pipeline and performed the data analysis. M. Tewari, S.W.C and M.Tuck designed the human subjects studies. M. Tuck and A.G. carried out human subjects studies and A.G. performed clinical data abstraction. All authors read the manuscript and provided input.

## Competing Interests

The authors declare no competing financial interests.

## Appendix

**Table S1**. Sequences of the equimolar synthetic pool.

**Table S2**. Demographic data for healthy control participants included in the study.

**Table S3**. mRNA and lncRNA fragments detected in plasma from healthy individuals using phospho-sRNA-seq (PNK) and standard small RNA-seq (No.PNK) combined with a custom analysis pipeline.

**Table S4**. Table of co-expression clustering results from dynamically-expressed genes in bone marrow transplant patients, P04 and P07.

**Table S5**. Databases used for aligning contaminants.

## References

Adiconis X, Borges-Rivera D, Satija R, DeLuca DS, Busby MA, Berlin AM, Sivachenko A, Thompson DA, Wysoker A, Fennell T, Gnirke A, Pochet N, Regev A & Levin JZ (2013) Comparative analysis of RNA sequencing methods for degraded or low-input samples. Nat. Methods 10: 623–629

Arroyo JD, Chevillet JR, Kroh EM, Ruf IK, Pritchard CC, Gibson DF, Mitchell PS, Bennett CF, Pogosova-Agadjanyan EL, Stirewalt DL, Tait JF & Tewari M (2011) Argonaute2 complexes carry a population of circulating microRNAs independent of vesicles in human plasma. Proc. Natl. Acad. Sci. U. S. A. 108: 5003–5008

Ben-Porath I, Thomson MW, Carey VJ, Ge R, Bell GW, Regev A & Weinberg RA (2008) An embryonic stem cell-like gene expression signature in poorly differentiated aggressive human tumors. Nat. Genet. 40: 499–507

Cameron V & Uhlenbeck OC (1977) 3′-Phosphatase activity in T4 polynucleotide kinase. Biochemistry 16: 5120–5126

Cheng HH, Yi HS, Kim Y, Kroh EM, Chien JW, Eaton KD, Goodman MT, Tait JF, Tewari M & Pritchard CC (2013) Plasma processing conditions substantially influence circulating microRNA biomarker levels. PLoS One 8: e64795

Chen H-Y, Yu S-L, Chen C-H, Chang G-C, Chen C-Y, Yuan A, Cheng C-L, Wang C-H, Terng H-J, Kao S-F, Chan W-K, Li H-N, Liu C-C, Singh S, Chen WJ, Chen JJW & Yang P-C (2007) A five-gene signature and clinical outcome in non-small-cell lung cancer. N. Engl. J. Med. 356: 11–20

Chen L, Reeve J, Zhang L, Huang S, Wang X & Chen J (2018) GMPR: A robust normalization method for zero-inflated count data with application to microbiome sequencing data. PeerJ 6: e4600

Chung D, Kuan PF, Li B, Sanalkumar R, Liang K, Bresnick EH, Dewey C & Keleş S (2011) Discovering transcription factor binding sites in highly repetitive regions of genomes with multi-read analysis of ChIP-Seq data. PLoS Comput. Biol. 7: e1002111

Cuchillo CM, Nogués MV & Raines RT (2011) Bovine Pancreatic Ribonuclease: Fifty Years of the First Enzymatic Reaction Mechanism. Biochemistry 50: 7835–7841

Danielson KM, Rubio R, Abderazzaq F, Das S & Wang YE (2017) High Throughput Sequencing of Extracellular RNA from Human Plasma. PLoS One 12: e0164644

Dobin A, Davis CA, Schlesinger F, Drenkow J, Zaleski C, Jha S, Batut P, Chaisson M & Gingeras TR (2013) STAR: ultrafast universal RNA-seq aligner. Bioinformatics 29: 15–21

Freedman JE, Gerstein M, Mick E, Rozowsky J, Levy D, Kitchen R, Das S, Shah R, Danielson K, Beaulieu L, Navarro FCP, Wang Y, Galeev TR, Holman A, Kwong RY, Murthy V, Tanriverdi SE, Koupenova-Zamor M, Mikhalev E & Tanriverdi K (2016) Diverse human extracellular RNAs are widely detected in human plasma. Nat. Commun. 7: 11106

Giraldez MD, Spengler RM, Etheridge A, Godoy PM, Barczak AJ, Srinivasan S, De Hoff PL, Tanriverdi K, Courtright A, Lu S, Khoory J, Rubio R, Baxter D, Driedonks TAP, Buermans HPJ, Nolte-’t Hoen ENM, Jiang H, Wang K, Ghiran I, Wang YE, et al (2018) Comprehensive multi-center assessment of small RNA-seq methods for quantitative miRNA profiling. Nat. Biotechnol. 36: 746–757

Godoy PM, Bhakta NR, Barczak AJ, Cakmak H, Fisher S, MacKenzie TC, Patel T, Price RW, Smith JF, Woodruff PG & Erle DJ (2018) Large Differences in Small RNA Composition Between Human Biofluids. Cell Rep. 25: 1346–1358

Hafner M, Landgraf P, Ludwig J, Rice A, Ojo T, Lin C, Holoch D, Lim C & Tuschl T (2008) Identification of microRNAs and other small regulatory RNAs using cDNA library sequencing. Methods 44: 3–12

Ha M & Kim VN (2014) Regulation of microRNA biogenesis. Nat. Rev. Mol. Cell Biol. 15: 509–524

Huang X, Yuan T, Tschannen M, Sun Z, Jacob H, Du M, Liang M, Dittmar RL, Liu Y, Liang M, Kohli M, Thibodeau SN, Boardman L & Wang L (2013) Characterization of human plasma-derived exosomal RNAs by deep sequencing. BMC Genomics 14: 319

Hu G, Drescher KM & Chen X-M (2012) Exosomal miRNAs: Biological Properties and Therapeutic Potential. Front. Genet. 3: 56

Hunter MP, Ismail N, Zhang X, Aguda BD, Lee EJ, Yu L, Xiao T, Schafer J, Lee M-LT, Schmittgen TD, Nana-Sinkam SP, Jarjoura D & Marsh CB (2008) Detection of microRNA expression in human peripheral blood microvesicles. PLoS One 3: e3694

Iyer MK, Niknafs YS, Malik R, Singhal U, Sahu A, Hosono Y, Barrette TR, Prensner JR, Evans JR, Zhao S, Poliakov A, Cao X, Dhanasekaran SM, Wu Y-M, Robinson DR, Beer DG, Feng FY, Iyer HK & Chinnaiyan AM (2015) The landscape of long noncoding RNAs in the human transcriptome. Nat. Genet. 47: 199–208

Jain A & Tuteja G (2018) TissueEnrich: Tissue-specific gene enrichment analysis. Bioinformatics Available at: http://dx.doi.org/10.1093/bioinformatics/bty890

Kamm RC & Smith AG (1972) Ribonuclease activity in human plasma. Clin. Biochem. 5: 198–200

Knight SW & Bass BL (2001) A role for the RNase III enzyme DCR-1 in RNA interference and germ line development in Caenorhabditis elegans. Science 293: 2269–2271

Koh W, Pan W, Gawad C, Fan HC, Kerchner GA, Wyss-Coray T, Blumenfeld YJ, El-Sayed YY & Quake SR (2014) Noninvasive in vivo monitoring of tissue-specific global gene expression in humans. Proc. Natl. Acad. Sci. U. S. A. 111: 7361–7366

Lamm AT, Stadler MR, Zhang H, Gent JI & Fire AZ (2011) Multimodal RNA-seq using single-strand, double-strand, and CircLigase-based capture yields a refined and extended description of the C. elegans transcriptome. Genome Res. 21: 265–275

Langfelder P & Horvath S (2008) WGCNA: an R package for weighted correlation network analysis. BMC Bioinformatics 9: 559

Lee C, Harris RA, Wall JK, Mayfield RD & Wilke CO (2013) RNaseIII and T4 polynucleotide Kinase sequence biases and solutions during RNA-seq library construction. Biol. Direct 8: 16

Lee Y, Ahn C, Han J, Choi H, Kim J, Yim J, Lee J, Provost P, Rådmark O, Kim S & Kim VN (2003) The nuclear RNase III Drosha initiates microRNA processing. Nature 425: 415–419

Leng N, Li Y, McIntosh BE, Nguyen BK, Duffin B, Tian S, Thomson JA, Dewey CN, Stewart R & Kendziorski C (2015) EBSeq-HMM: a Bayesian approach for identifying gene-expression changes in ordered RNA-seq experiments. Bioinformatics 31: 2614–2622

Liu X, Yu X, Zack DJ, Zhu H & Qian J (2008a) TiGER: a database for tissue-specific gene expression and regulation. BMC Bioinformatics 9: 271

Liu X, Yu X, Zack DJ, Zhu H & Qian J (2008b) TiGER: a database for tissue-specific gene expression and regulation. BMC Bioinformatics 9: 271

Ludwig N, Leidinger P, Becker K, Backes C, Fehlmann T, Pallasch C, Rheinheimer S, Meder B, Stähler C, Meese E & Keller A (2016) Distribution of miRNA expression across human tissues. Nucleic Acids Res. 44: 3865–3877

Lu L, Li J, Moussaoui M & Boix E (2018) Immune Modulation by Human Secreted RNases at the Extracellular Space. Front. Immunol. 9: 1012

Martin M (2011) Cutadapt removes adapter sequences from high-throughput sequencing reads. EMBnet.journal 17: 10–12

Max KEA, Bertram K, Akat KM, Bogardus KA, Li J, Morozov P, Ben-Dov IZ, Li X, Weiss ZR, Azizian A, Sopeyin A, Diacovo TG, Adamidi C, Williams Z & Tuschl T (2018) Human plasma and serum extracellular small RNA reference profiles and their clinical utility. Proc. Natl. Acad. Sci. U. S. A. 115: E5334–E5343

Mortazavi A, Williams BA, McCue K, Schaeffer L & Wold B (2008) Mapping and quantifying mammalian transcriptomes by RNA-Seq. Nat. Methods 5: 621–628

Novogrodsky A & Hurwitz J (1966) The enzymatic rylation of ribonucleic acid and deoxyribonucleic acid. I. Phosphorylation at 5′-hydroxyl termini. J. Biol. Chem. 241: 2923–2932

Perou CM, Sørlie T, Eisen MB, van de Rijn M, Jeffrey SS, Rees CA, Pollack JR, Ross DT, Johnsen H, Akslen LA, Fluge O, Pergamenschikov A, Williams C, Zhu SX, Lønning PE, Børresen-Dale AL, Brown PO & Botstein D (2000) Molecular portraits of human breast tumours. Nature 406: 747–752

Potti A, Mukherjee S, Petersen R, Dressman HK, Bild A, Koontz J, Kratzke R, Watson MA, Kelley M, Ginsburg GS, West M, Harpole DH & Nevins JR (2006) A Genomic Strategy to Refine Prognosis in Early-Stage Non–Small-Cell Lung Cancer. N. Engl. J. Med. 355: 570–580

Richardson CC (1965) Phosphorylation of nucleic acid by an enzyme from T4 bacteriophage-infected Escherichia coli. Proc. Natl. Acad. Sci. U. S. A. 54: 158–165

Roser AE, Caldi Gomes L, Schünemann J, Maass F & Lingor P (2018) Circulating miRNAs as Diagnostic Biomarkers for Parkinson’s Disease. Front. Neurosci. 12: 625

Schwarzenbach H, Hoon DSB & Pantel K (2011) Cell-free nucleic acids as biomarkers in cancer patients. Nat. Rev. Cancer 11: 426–437

Shah MY & Calin GA (2014) MicroRNAs as therapeutic targets in human cancers. Wiley Interdiscip. Rev. RNA 5: 537–548

Turchinovich A, Surowy H, Serva A, Zapatka M, Lichter P & Burwinkel B (2014) Capture and Amplification by Tailing and Switching (CATS) An ultrasensitive ligation-independent method for generation of DNA libraries for deep sequencing from picogram amounts of DNA and RNA. RNA Biol. 11: 817–828

Vvedenskaya IO, Goldman SR & Nickels BE (2015) Preparation of cDNA libraries for high-throughput RNA sequencing analysis of RNA 5′ ends. Methods Mol. Biol. 1276: 211–228

Wang Z, Gerstein M & Snyder M (2009) RNA-Seq: a revolutionary tool for transcriptomics. Nat. Rev. Genet. 10: 57–63

Wu X, Kim T-K, Baxter D, Scherler K, Gordon A, Fong O, Etheridge A, Galas DJ & Wang K (2017) sRNAnalyzer-a flexible and customizable small RNA sequencing data analysis pipeline. Nucleic Acids Res. 45: 12140–12151

Yeri A, Courtright A, Reiman R, Carlson E, Beecroft T, Janss A, Siniard A, Richholt R, Balak C, Rozowsky J, Kitchen R, Hutchins E, Winarta J, McCoy R, Anastasi M, Kim S, Huentelman M & Van Keuren-Jensen K (2017) Total Extracellular Small RNA Profiles from Plasma, Saliva, and Urine of Healthy Subjects. Sci. Rep. 7: 44061

Yuan T, Huang X, Woodcock M, Du M, Dittmar R, Wang Y, Tsai S, Kohli M, Boardman L, Patel T & Wang L (2016) Plasma extracellular RNA profiles in healthy and cancer patients. Sci. Rep. 6: 19413

Zhang Y, Liu D, Chen X, Li J, Li L, Bian Z, Sun F, Lu J, Yin Y, Cai X, Sun Q, Wang K, Ba Y, Wang Q, Wang D, Yang J, Liu P, Xu T, Yan Q, Zhang J, et al (2010) Secreted monocytic miR-150 enhances targeted endothelial cell migration. Mol. Cell 39: 133–144

